# Chronic Hypoxic Signalling Reprograms Metabolism and Alters Redox and Lipid Homeostasis in Rod Photoreceptors

**DOI:** 10.64898/2026.07.03.736307

**Authors:** Larissa P. Govers, Daniel T. Hass, Martin-Paul Agbaga, Claudia Matter, Antonia Fottner, Marijana Samardzija, James B. Hurley, Christian Grimm

## Abstract

Photoreceptors are among the most metabolically active cells in the retina and are therefore highly sensitive to fluctuations in oxygen availability. Age-related tissue changes in the eye affect oxygen delivery to the outer retina, which may result in hypoxic stress within photoreceptors and can contribute to disease development and retinal degeneration. To investigate how chronic hypoxic signalling affects photoreceptor metabolism, we examined a rod-specific *Vhl* knockout mouse (Rod^Δ*Vhl*^), in which constitutive HIF activation mimics the molecular response to hypoxia. Combining a cell-type–enriched multi-omics approach with metabolic flux analysis, we identified an early metabolic response in the retina of Rod^Δ*Vhl*^ mice prior to degeneration. This response was characterized by a shift towards an oxidative redox environment indicated by a decrease in nucleotide precursors and an increased antioxidant response. While steady-state glycolytic flux remained unchanged, the dynamic ^13^C-glucose tracing revealed accelerated carbon flow through the three-carbon glycolytic intermediates, indicating a carbon rerouting. Outer segment lipidomics revealed selective remodelling of phosphatidylcholine and phosphatidylethanolamine species toward more oxidation-resistant and elongated acyl chains, supported by early gene upregulation of essential enzymes involved in fatty acid elongation, desaturation and oxidation. Together, these findings indicate a coordinated shift in metabolic and lipid pathways in photoreceptors under chronic hypoxic stress, consistent with an adaptive response that may help preserve outer segment integrity and improve stress resilience.

## Introduction

The retina has one of the highest metabolic demands in the human body [1]. Within the retina, rod and cone photoreceptors possess the highest density of mitochondria, making them highly dependent on efficient delivery of oxygen, glucose and other nutrients from the choroidal vasculature [2]. Otto Warburg found that the metabolism of these nutrients is unique in the retina. Rather than efficient oxidation of glucose via glycolysis coupled to oxidative phosphorylation (OXPHOS), retinas will ferment glucose to lactate even when oxygen is present [3]. This process is known as aerobic glycolysis or the Warburg effect. Using fluorescence lifetime imaging of metabolic biosensors, we recently demonstrated that photoreceptors rely on both OXPHOS and glycolysis to fully sustain their intracellular ATP levels [4]. This dual dependency satisfies both the energetic and anabolic demands of photoreceptors, with glycolysis providing the intermediates needed for outer segment biogenesis and the daily renewal of its outermost tip [5].

Aging impairs choroidal blood flow, thereby disturbing oxygen delivery to the retinal pigment epithelium (RPE) and outer retina [6, 7]. As a consequence, photoreceptors may experience a hypoxic environment that imposes metabolic stress [8]. Changes in photoreceptor metabolism have been suggested to be at the core of several blinding diseases, including age-related macular degeneration [9, 10]. Thus, hypoxia not only alters cellular metabolism but may also contribute to the induction and/or progression of retinal pathologies, making the understanding of its consequences for retinal cells mandatory for the development of successful therapies [8].

The limited oxygen supply during hypoxia not only alters the mitochondrial redox state and impairs OXPHOS but also activates the hypoxia-inducible transcription factor (HIF) pathway, which regulates the cellular adaptation to reduced oxygen availability in affected cells. While the former primarily affects mitochondrial metabolism, the latter dictates how cells adjust to a hypoxic environment and is often activated in disease states, where it is known to drive metabolic changes [11]. The HIF pathway is regulated, in part, by the von Hippel-Lindau tumor-suppressor protein (VHL). A rod-specific knockout of the *Vhl* gene (Rod^Δ*Vhl*^) results in chronic HIF activation, leading to late-onset, slowly progressing photoreceptor degeneration [8, 12, 13]. The absence of VHL increases lactate production in rod photoreceptors, accompanied by a severe downregulation of proteins involved in both OXPHOS and the citric acid (TCA) cycle. When only OXPHOS was downregulated, the degeneration was delayed and less severe compared to the degeneration in Rod^Δ*Vhl*^ mice [13]. This suggests that while the high glycolytic rate that is intrinsic to rods can generate sufficient ATP to maintain the function of the cell for some time, a chronic hypoxic response may introduce additional metabolic stressors that compromise the anabolic pathways required for processes such as outer segment renewal, ultimately leading to degeneration.

In this study, we addressed the metabolic changes in photoreceptors in a mouse model subjected to rod-specific chronic pseudo-hypoxia before the onset of degeneration, with the aim of identifying how sustained HIF activation shapes photoreceptor metabolism and cellular adaptations.

## Methods

### Animals

All animal experiments were approved by the Cantonal Veterinary Office of Zurich, Switzerland (license numbers: ZH199/2022, ZH069/2025) and adhered to the ARVO statement for the use of Animals in Ophthalmic and Vision Research. Mice were maintained as breeding colonies at the Laboratory Animal Services Center (LASC) of the University of Zurich with a 14/10 h light-dark cycle with an average light intensity of 60-150 lux at cage level, depending on the position on the rack. Mice had access to food and water *ad libitum*.

Generation of Rod^Δ*Vhl*^ (*Vhl*^flox/flox^;Opsin-Cre) mice by intercrossing *Vhl*^flox/flox^ and Opsin-cre (LMPOC1) mice was described previously [14]. Breeding pairs consisted of hemizygous Opsin-Cre mice and Cre-negative mating partners, giving rise to Rod^Δ*Vhl*^ and *Vhl*^flox/flox^ mice that served as controls. Cre expression in rods in Opsin-Cre mice starts around postnatal day 7 and increases until 6 weeks of age [14]. All mice were homozygous for the *Rpe65_450Leu_* variant. Mice were euthanized with CO_2_ followed by decapitation.

### Proteomics analysis

Previously published proteomics data of individual retinal layers [15] was newly analyzed after excluding proteins involved in OXPHOS and glycolysis [13]. Functional annotation of the remaining proteins was performed using Gene Ontology resources for the photoreceptor segments and outer nuclear layer (p≤0.05) [16, 17]. Normalized protein abundance ratios were visualized with GraphPad Prism (GraphPad Software, San Diego, CA, USA).

### Metabolomics, sample preparation

Samples were prepared using the ReLayS method [15]. In short, retinas from 10-week-old Rod^Δ*Vhl*^ and control mice (N=12 per genotype, equal distribution of males and females) were isolated and placed in PBS on ice. Retinas were cleaned from vitreous and halved through the optic nerve head. After flattening on a polyvinylidene difluoride (PVDF) membrane with the photoreceptors facing up, the half-retinas were frozen on the membrane using a metal platform on dry ice. Photoreceptor segments and the outer nuclear layer were separated based on their adherence to another PVDF membrane. After physical separation, the tissue on the membranes was frozen again on a metal platform on dry ice and stored at -80°C until further use. To isolate polar metabolites, tissue samples on the membranes were homogenized by sonication with an ultrasonic homogenizer in an ice-cold solution containing acetonitrile:methanol:water (40:40:20, v/v/v). After sonication, samples were incubated at -20°C for 1 h, centrifuged at maximum speed for 10 min and the supernatants collected. The protein pellet was dissolved in ice-cold Tris-HCl (100 mM, pH 8.0) containing protease inhibitors (P2714, Sigma-Aldrich, St. Louis, MO, USA) and the protein concentration was determined with a BCA assay (Thermo Fisher Scientific, Waltham, MA, USA).

Prior to measuring, extracts were dried under nitrogen flow and reconstituted in injection buffer (90% acetonitrile). The solution was vortexed and centrifuged for 10 min at 12000 rpm and 4°C. The clear supernatants were transferred to glass vials with narrowed bottoms (Total Recovery Vials, Waters) for LC-MS injection. In addition, method blanks, QC standards, and pooled samples were prepared in the same way to serve as quality control for the measurements.

### Metabolomics, LC-MS analysis

Untargeted metabolomics was performed at the Functional Genomics Center Zurich (FGCZ). Polar metabolites were separated on a Thermo Vanquish Horizon Binary Pump equipped with a Waters Premier BEH Amide column (150 mm x 2.1 mm), applying a gradient of 10 mM ammonium bicarbonate in water (A) and 10 mM ammonium bicarbonate in 95% acetonitrile (B) from 99% B to 30% B over 12 min. The injection volume was 5 μL. The flow rate was 0.4 μL/min with a column temperature of 40°C and an autosampler temperature of 5°C. The LC was coupled to Thermo Q Exactive mass spectrometer by a HESI source. MS1 (molecular ion) and MS2 (fragment) data were acquired using negative polarization and Full MS / dd-MS² (Top5) over a mass range of 70 to 1050 m/z at MS1 resolution of 60’000 and MS2 resolution of 7’500.

### Metabolomics, untargeted data analysis

The metabolomics data set was evaluated in an untargeted fashion with Compound Discoverer software (Thermo Scientific). The modular data analysis workflow included spectra selection, retention times alignment, compound detection and grouping, gap filling, background filtering and normalization (data are protein content normalized). mzCloud and mzVault have been used to score fragmentation patterns and assign MS2-based identities to the features. A filtering process was performed, leading to the manually annotated compound table, where each feature is annotated with the highest level of confidence. Filtering parameters used were the following: Signal-to-noise ratio > 3, mzCloud or mzVault match >50, ppm mass error within +/- 5 ppm, match with in-house developed MS1_RT library within +/-10 s, chromatographic peak, and MS2 spectra quality. Quality controls were run on pooled QC samples and reference compound mixtures to determine technical accuracy and stability.

Data was analyzed using MetaboAnalyst (Version 6.0) [18]. Peak intensities were visualized with Prism software.

### Lipid profiling

Retinas from 10-week-old Rod^Δ*Vhl*^ and control mice (N=6 samples per genotype; 4 retinas from 2 mice per sample; equal distribution of males and females) were isolated using the Winkler method and stored at -80°C [19]. Retinal outer segment membranes were prepared according to published protocols using discontinuous sucrose gradient ultracentrifugation [20]. The lipid profiling was then performed as previously described [9]. In short, tissue samples were homogenized in 40% methanol and subsequently diluted 1:40 with a 2-propanol/methanol/chloroform mixture (4:2:1, v/v/v) containing 20 mM ammonium formate and internal standards from Avanti Research (Alabaster, AL, USA): 1.0 μM phosphatidylcholine (PC) 14:0/14:0 (Cat #850345), 1.0 μM phosphatidylethanolamine (PE) 14:0/14:0 (Cat #850745), and 0.33 μM phosphatidylserine (PS) 14:0/14:0 (Cat #840033). Samples were analyzed by direct infusion using a chip-based nano-electrospray ionization source (Advion NanoMate; Advion Interchim, Ithaca, NY, USA) coupled to a triple-quadrupole mass spectrometer (TSQ Ultra, Thermo Fisher Scientific). PC species were quantified using precursor ion scanning of m/z 184, while PE and PS species were analyzed by neutral loss scanning of m/z 141 and m/z 185, respectively. Lipid species were expressed as relative percentages of the total signal based on response values, and molecular species abundances were calculated using Lipid Mass Spectrum Analysis (LIMSA) software (University of Helsinki, Finland).

### Aconitase activity

Aconitase activity was determined using the Aconitase Activity Assay Kit (ab109712, Abcam, Cambridge, UK). In short, whole retinas from Rod^Δ*Vhl*^ and control mice were isolated at 10 weeks of age and mechanically homogenized in aconitase preservation solution using a 21G needle. The samples were pipetted into a 96-well plate, and activity buffer with 1/25 volume isocitrate and 1/100 volume manganese solution from the kit was added. The absorbance at 240 nm was measured at one-minute intervals for 60 min using the BioTek Synergy H1 Multimode Reader (Agilent, Santa Clara, CA, USA). The maximum velocity (V_max_) per retina was plotted and visualized using Prism software.

### Glutathione S-Transferase assay

Glutathione S-Transferase (GST) activity was measured using the GST Assay Kit (CS0410, Sigma-Aldrich). Whole retinas from Rod^Δ*Vhl*^ and control mice were isolated at 10 weeks of age, mechanically homogenized in sample buffer provided by the kit and loaded into a 96-well plate. A master mix containing 2 mM L-glutathione and 1 mM 1-chloro-2,4-dinitrobenzene (CDNB) was added to each well. Absorbance was measured at 340 nm every minute for 60 min using the BioTek Synergy H1 plate reader. Change in absorbance was calculated over time and normalized to protein content that was determined with a BCA assay (Thermo Fisher Scientific).

### *Ex vivo* glucose tracing

Retinas were isolated and placed in ice-cold Hank’s buffered salt solution (Thermo Fisher Scientific) to remove the vitreous and any other tissue contaminants. The retinas were then incubated in Krebs-Ringer Bicarbonate (KRB) buffer supplemented with 5 mM [U-^13^C] glucose for 90 min at 37°C to reach isotopic steady-state. The medium was sampled at 0, 30, 60, and 90 min. To study dynamic metabolic flux, isolated retinas and eyecups were incubated in KRB buffer supplemented with 5 mM [U-^13^C] glucose for 0, 1, and 5 min. Eyecup measurements had an additional time point at 3 minutes. After washing in Hank’s buffered salt solution, the tissue samples were snap frozen in liquid nitrogen for analysis. Metabolites were extracted in 80% methanol, derivatized for GC-MS and LC-MS, and quantified as described previously [21].

### Semi-quantitative real-time PCR

Whole retinas from Rod^Δ*Vhl*^ and control mice were isolated at 10 weeks, 4 months and 6 months of age. Total RNA was isolated from the tissue using an RNA isolation kit (NucleoSpin RNA, Macherey-Nagel GmbH & co.KG, Duren, Germany) according to the manufacturer’s instructions, including an on-column DNase digestion step (740963, Macherey-Nagel GmbH & co.KG). cDNA synthesis was performed using 650 ng total RNA, 20 pmol of oligo-(d)T primers and M-MLV reverse transcriptase (Promega, Dubendorf, Switzerland). Gene expression was measured by real-time PCR using 10 ng of cDNA template, PowerUP SYBR Green Master Mix (Thermo Fisher Scientific) and specific primer pairs (*Actb* fwd 5’-3’: CAACGGCTCCGGCATGTGC and rev 5’-3’: CTCTTGCTCTGGGCCTCG*; Scd2* fwd 5’-3’: TCATAGCACCCTTTGTCGCT and rev 5’-3’: GATTGTGGTGGTGGCTGAGTA; *Acaca* fwd 5’-3’: TGTCACCAGCCTCCGTC and rev 5’-3’: GTGAAATCTCGTTGTGAGTCTA*; Cpt1a* fwd 5’-3’: TGATGACGGCTATGGTGT and rev 5’-3’: GCGGTGTGAGTCTGTCT). The analysis was performed with the ABI QuantStudio 3 system (Thermo Fisher Scientific). *Actb* was used as the housekeeping gene. Relative gene expression was calculated using the 2^-ddCt^ method [22] and visualized with Prism software.

### Statistical analysis

Statistical analysis was performed using Prism software. Significance was tested by an unpaired t-test when two independent groups were compared. For comparisons across multiple time points, an ordinary two-way ANOVA followed by Tukey’s multiple comparison test was applied, while lipid profiling data were analyzed using an ordinary two-way ANOVA with Sidak’s multiple comparison test. Data were considered significantly different with p ≤ 0.05. All numbers of animals/samples are indicated in the figure legends.

## Results

### Chronic HIF activation establishes an oxidative redox environment in photoreceptors

Rod photoreceptors rely on both glycolysis and OXPHOS to meet their metabolic demand [4, 5]. Previous work has demonstrated that rod photoreceptors with chronically active HIF transcription factors (Rod^Δ*Vhl*^) undergo a metabolic shift from OXPHOS to increased glycolysis, which precedes, and potentially contributes to, late-onset and progressive photoreceptor degeneration [13]. Within that work, proteomics data on separated layers of the retina revealed a reduction of OXPHOS proteins as well as an increase in enzymes involved in early glycolysis in photoreceptor segments and the outer nuclear layer [13]. To identify potential changes in metabolic pathways beyond primary ATP-generating processes, we reanalyzed the published proteomics dataset for the photoreceptor inner/outer segments and the outer nuclear layer after *in silico* removal of all proteins annotated to OXPHOS and glycolysis (Figure 1A). This allowed us to uncover additional metabolic adaptations before the onset of degeneration that were otherwise masked by the dominant OXPHOS/glycolysis alterations.

**Figure 1.**
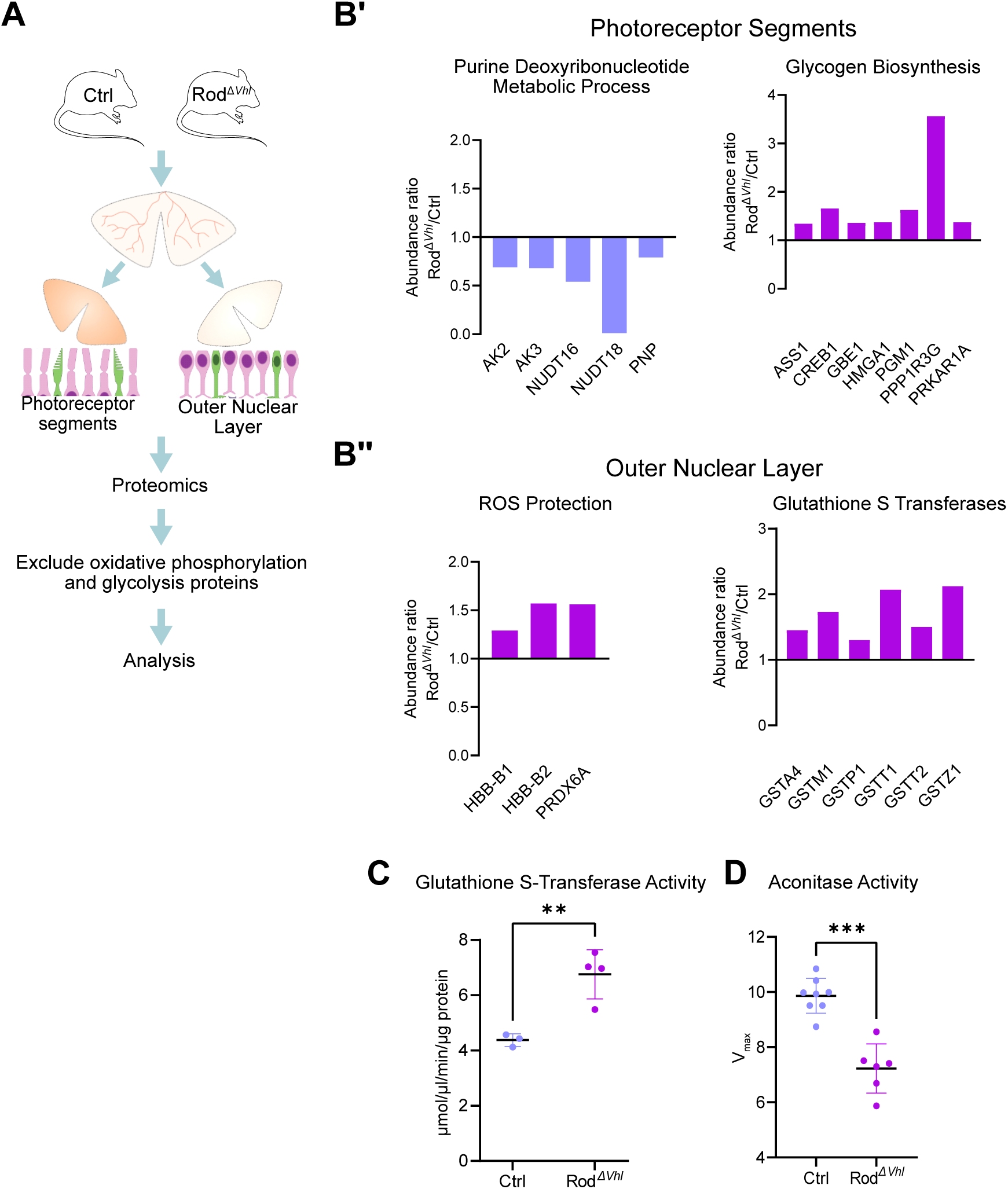
Reanalysis of proteomics data reveals an adapted redox status in Rod^Δ*Vhl*^ mice. A, Schematic of the proteomics workflow showing retinal layer separation and removal of glycolysis/OXPHOS-annotated proteins for focused pathway analysis. B’, Significantly altered proteins in the photoreceptor segment layer. B’’, Significantly altered proteins in the outer nuclear layer. Proteomics data for reanalysis were from [13]. C, Glutathione S-transferase activity normalized to protein content in isolated whole retina lysates of Rod^Δ*Vhl*^ and control mice at 10 weeks of age. D, Aconitase activity in lysates of entire retinas from Rod^Δ*Vhl*^ and control mice at 10 weeks of age. Shown are individual data points, mean ± SD. p ≤ 0.01. ***: p ≤ 0.001.

Reanalysis of the data (Table S1) revealed a decrease of proteins (adenylate kinase 2 (AK2), AK3; nudix hydrolase 16 (NUDT16); NUDT18 and purine nucleoside phosphorylase (PNP)) that are essential for nucleotide turnover and the salvage pathway and thus are involved in purine deoxyribonucleotide metabolism in the photoreceptor segments. Furthermore, several glycogen biosynthesis proteins were upregulated in the segments, indicating a metabolic stress response (Figure 1B’). The proteomics dataset of the outer nuclear layer showed an increase in hemoglobin subunit beta-B1 (HBB-B1) and HBB-B2 which are important for enhanced reactive oxygen species (ROS) buffering [23], as well as peroxiredoxin 6 (PRDX6A), which is a lipid-focused antioxidant [24]. Expression of these genes in photoreceptors has been documented by others in single-cell transcriptomics from mouse and human retinas [25–27]. Multiple glutathione S-transferases were upregulated on protein level (Figure 1B’’) indicating an activation of antioxidant pathways. To confirm this, we performed a glutathione S-transferase activity assay which indeed showed increased activity in retinas of Rod*^ΔVhl^* (Figure 1C). In parallel, we tested another indicator of oxidative stress, aconitase, which catalyzes the second step of the TCA and contains an Fe-S cluster that is highly sensitive to oxidation [28]. We found decreased aconitase activity (Figure 1D), which is consistent with the expected enzyme inactivation under conditions of elevated ROS. These proteomic and functional changes revealed a shift towards reduced nucleotide turnover, increased glycogen metabolism, as well as an increased antioxidant response, which indicates an altered redox state and metabolic stress adaptation in the affected photoreceptors.

Identifying the metabolic signature of photoreceptors during the chronic hypoxic response before degeneration would deepen our knowledge of the mechanisms that may lead to disease onset and progression. To get insight into these mechanisms, metabolomics was performed on tissue samples containing the photoreceptor segments and outer nuclear layer that were isolated from the whole retina using the ReLayS method (Figure 2A, Table S2). The heatmap of relative metabolite abundances across samples showed clustering of the Rod*^ΔVhl^* and control mice, indicating distinct metabolic profiles in the photoreceptors of the two genotypes (Figure 2B).

**Figure 2.**
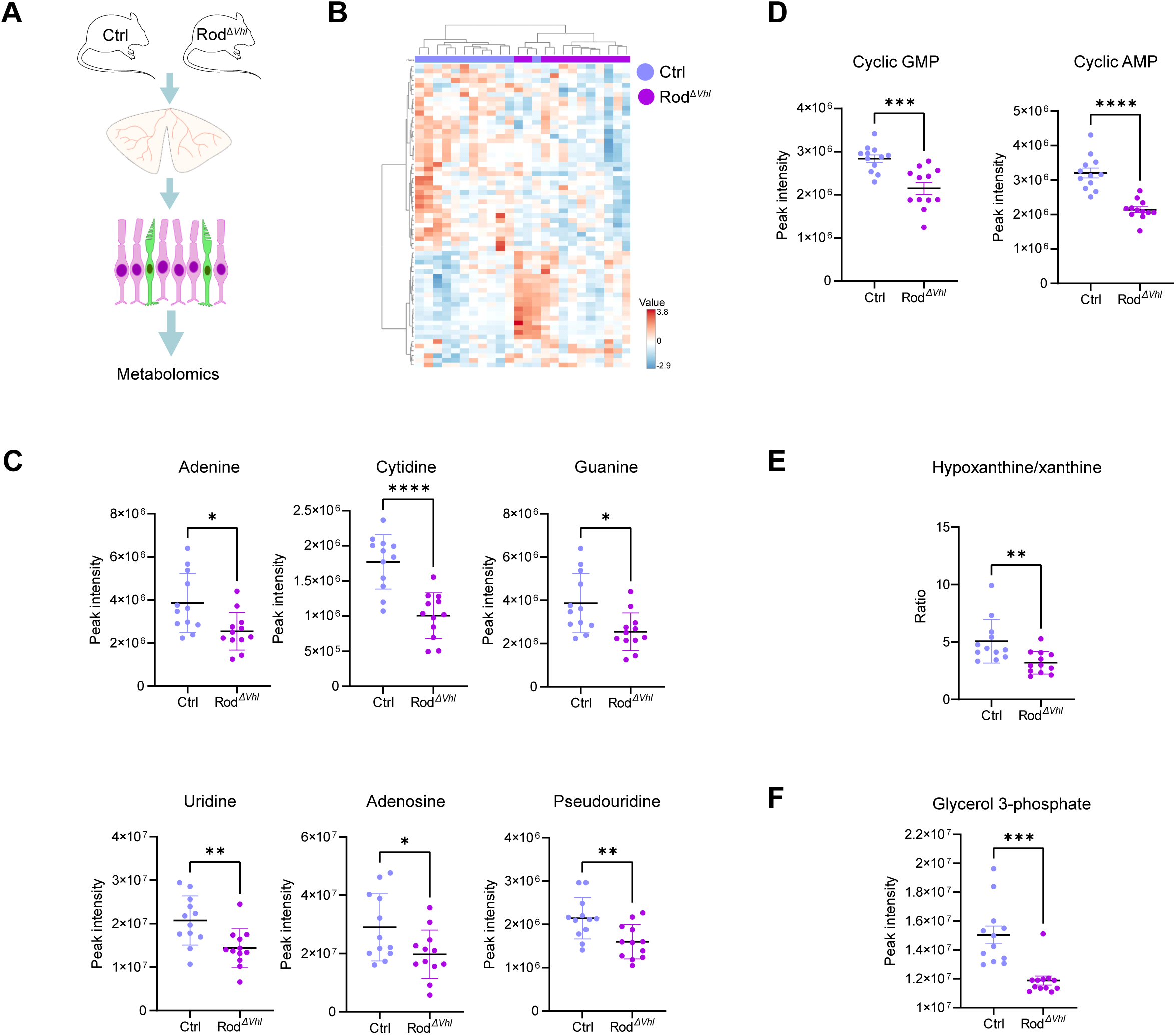
Metabolomic profiling reveals imbalance in nucleotide-related metabolites in the outer retina of Rod^Δ*Vhl*^ mice. A, Schematic of the metabolomics workflow showing separation of the photoreceptor layer (outer retina) followed by metabolite quantification in photoreceptors of 10-week-old Rod^Δ*Vhl*^ and control mice. B, Heatmap of significantly altered metabolites and clustering of samples from Rod^Δ*Vhl*^ and control mice. C, Nucleotide precursor metabolites in photoreceptors of Rod^Δ*Vhl*^ and control mice. D, Cyclic nucleotides cAMP and cGMP in photoreceptors of Rod^Δ*Vhl*^ and control mice. E, The ratio of hypoxanthine to xanthine in photoreceptors of Rod^Δ*Vhl*^ and control mice. F, The abundance of glycerol-3-phosphate in photoreceptors of Rod^Δ*Vhl*^ and control mice. N = 12 mice per genotype; both eyes of each mouse were pooled. Shown are individual data points, mean ± SD. *: p ≤ 0.05. **: p ≤ 0.01. ***: p ≤ 0.001. ****: p ≤ 0.0001.

Consistent with the decreased purine maintenance suggested by the proteomics dataset (Figure 1B’), nucleotide precursor metabolites (Figure 2C) and cyclic nucleotides cAMP and cGMP (Figure 2D) were significantly reduced, indicating reduced intracellular nucleotide pool availability. This was supported by the lower hypoxanthine/xanthine ratio in Rod*^ΔVhl^* mice (Figure 2E), which suggests reduced availability of purines resulting from decreased nucleotide pools and increased oxidative breakdown (Figure 2C). Increased oxidative stress might impact how photoreceptors handle metabolism and glucose. This is supported by the reduced level of glycerol-3-P (Figure 2F). Glycerol-3-P is normally produced during glycolysis, when glycerol-3-P dehydrogenase converts dihydroxyacetone phosphate (DHAP) into glycerol-3-P [29]. Indeed, both DHAP dynamics and the fatty acid signature were affected in the retina of Rod*^ΔVhl^* mice with their pseudo-hypoxic rod photoreceptors (see below).

Together, the proteomics and metabolomics datasets showed a consistent alteration of the metabolic profile in the retina of the Rod^Δ*Vhl*^ mice. The hypoxic response in photoreceptors was associated with decreased nucleotide-related metabolites and enhanced antioxidant defences, alongside altered glycolytic pathway dynamics, implying the presence of an oxidative redox environment and increased metabolic stress adaptation.

### Chronic HIF activation modulates glycolytic flux

While the omics-based approaches showed a distinct signature of affected molecular and metabolic pathways in the retina and photoreceptors of the Rod^Δ*Vhl*^ model, they were unable to capture the activity of the enzymes within these pathways. To address this limitation, isotope tracer studies using U-^13^C-glucose were performed on whole retinas that were extracted from Rod^Δ*Vhl*^ mice and their corresponding controls.

Isolated retinas were incubated in KRB buffer supplemented with U-^13^C-glucose, with the medium being sampled and analyzed at baseline, 30, 60 and 90 minutes to evaluate steady-state kinetics (Figure 3A). No significant differences were detected with respect to glucose uptake or lactate release by the retinas of Rod^Δ*Vhl*^ mice and their corresponding controls at any time point (Figure 3B). The total abundance of labelled metabolites in the retinal tissue also remained unchanged between genotypes at the incubation endpoint (90 minutes, Figure 3C). This combined data suggests that the overall glycolytic throughput was not altered at the level of the entire retina in Rod*^ΔVhl^* mice, despite the metabolic and redox alterations that were detected in the omics data.

**Figure 3.**
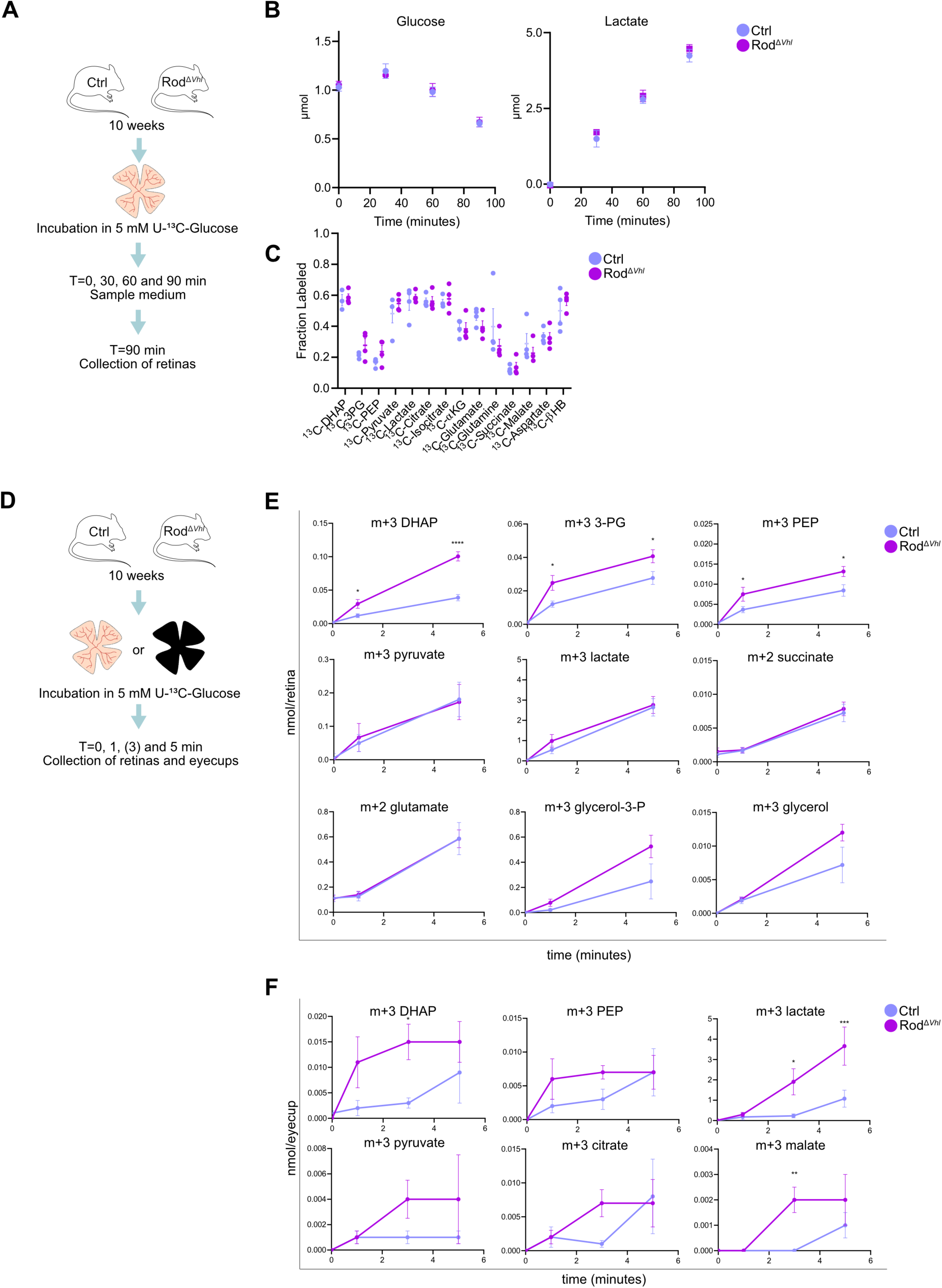
Glucose flux reveals altered intracellular carbon routing in Rod^Δ*Vhl*^ *ex vivo* retinas while preserving the overall glycolytic throughput. A, Schematic of the *ex vivo* ^13^C-glucose steady-state flux assay. Isolated retinas were incubated in medium containing 5 mM U-^13^C-glucose for 90 minutes, and medium or tissue was collected at the indicated time points. B, Steady-state flux measurements analysing glucose uptake or lactate release in media containing control and Rod^Δ*Vhl*^ retinal tissue over 90 min of incubation. (N = 4 mice (8 eyes) per genotype; left and right eyes of each mouse were pooled and analyzed together). C, Total abundance of ^13^C-labeled metabolites in control and Rod^Δ*Vhl*^ retinal tissue at 90 min of incubation. (N = 4 mice (8 eyes) per genotype; left and right eyes of each mouse were pooled and analyzed together). D, Schematic of the *ex vivo* ^13^C-glucose dynamic flux assay. Isolated retinas or eyecups (without retina) were incubated in KRB medium containing 5 mM U-^13^C-glucose for 0, 1, 3 (eyecups only) or 5 min. E, Incorporation of ^13^C in indicated metabolites in control and Rod^Δ*Vhl*^ retinal explants at 1-5 min. (N = 4 eyes (from 4 different mice) per genotype and timepoint). F. Incorporation of ^13^C in indicated metabolites in control and Rod^Δ*Vhl*^ eyecup tissue at 1–5 min. N = 4 eyes (from 4 different mice) per genotype and timepoint. Shown are individual data points, mean ± SEM. *: p ≤ 0.05. **: p ≤ 0.01. ****: p ≤ 0.0001.

Since steady-state glucose uptake and total ^13^C-label incorporation were unchanged, we performed dynamic flux analysis to understand how fast labelled carbons from glucose moved through the different metabolic enzymatic reactions over time. To study this, isolated retinas from Rod^Δ*Vhl*^ mice and controls were incubated in U-^13^C-glucose and sampled at 0, 1 and 5 minutes (Figure 3D). This showed a significantly faster labelling and thus increased flux of glucose-derived carbons to multiple metabolites, including DHAP, 3-phosphoglycerate (3-PG) and phosphoenolpyruvate (PEP) in the Rod*^ΔVhl^* retinas with their pseudo-hypoxic rods (Figure 3E). Furthermore, glycerol-3-P and glycerol showed a trend toward increased label incorporation, although this difference did not reach statistical significance. Metabolites such as pyruvate, lactate, succinate and glutamate were unaltered in Rod^Δ*Vhl*^ retinas (Figure 3E) indicating that the rate of lactate formation and the entry of the TCA cycle were similar in both genotypes under these conditions. This strongly suggests that chronic HIF activation in rod photoreceptors altered glucose shuttling, while preserving the terminal glycolytic and mitochondrial outputs. This observation is consistent with previously published work showing that chronic HIF activation does not enhance glycolytic flux but instead causes a metabolic shift [13].

Although the lack of VHL, and thus the hypoxic response in Rod*^ΔVhl^* mice, is specific to rods, dynamic flux analysis in the RPE-choroid tissue of Rod*^ΔVhl^* mice revealed trends of increased early and partially transient ^13^C incorporation into several metabolites, including DHAP, PEP, lactate, pyruvate, citrate and malate (Figure 3F). These differences suggest a secondary response of the RPE to the altered photoreceptor metabolism in Rod*^ΔVhl^* mice and further support the presence of a strong metabolic crosstalk between the RPE and photoreceptors. The rod-derived signal that may be altering RPE metabolism is not known.

### Chronic HIF activation drives selective phospholipid remodelling

As metabolic flux experiments revealed increased flux through the lower half of glycolysis, reflected by elevated labelling of DHAP, 3-PG and PEP, we investigated whether these changes were associated with alterations in lipid composition by performing shotgun-lipidomics on retinal outer segment preparations. The data revealed selective alterations in the phospholipid composition in the photoreceptor outer segments of Rod^Δ*Vhl*^ mice (Figure 4A). Phosphatidylcholine (PC) species containing monounsaturated fatty acids, including PC 34:1 and PC 36:1, were moderately increased, whereas the fully saturated PC 32:0 species were slightly reduced. In parallel, phosphatidylethanolamine (PE) species showed selective remodelling, with an increased abundance of the elongated, DHA-rich PE 40:6 species and a decrease in PE 38:6. In contrast, phosphatidylserine (PS) species remained unchanged. As the overall composition of major phospholipid classes (PC, PE, and PS) was largely preserved, these changes likely reflect selective lipid remodelling rather than a global disruption of phospholipid homeostasis. These lipid changes were supported by qPCR analysis of gene expression on the whole retina. Genes involved in fatty acid elongation (acetyl-CoA carboxylase 1; *Acaca)*, monounsaturated fatty acid production (stearoyl-CoA desaturase-2; *Scd2)*, and mitochondrial beta-oxidation (carnitine palmitoyltransferase 1A; *Cpt1a)* were upregulated in 10-week-old Rod*^ΔVhl^* mice (Figure 4B). This difference in gene expression was no longer detected after the onset of photoreceptor degeneration in these mice. These findings indicate that chronic HIF activation initiates a targeted reprogramming of lipid metabolism in rod photoreceptors before the onset of degeneration. Such a shift may represent an early compensatory response aimed at maintaining cellular integrity, function, and survival.

**Figure 4.**
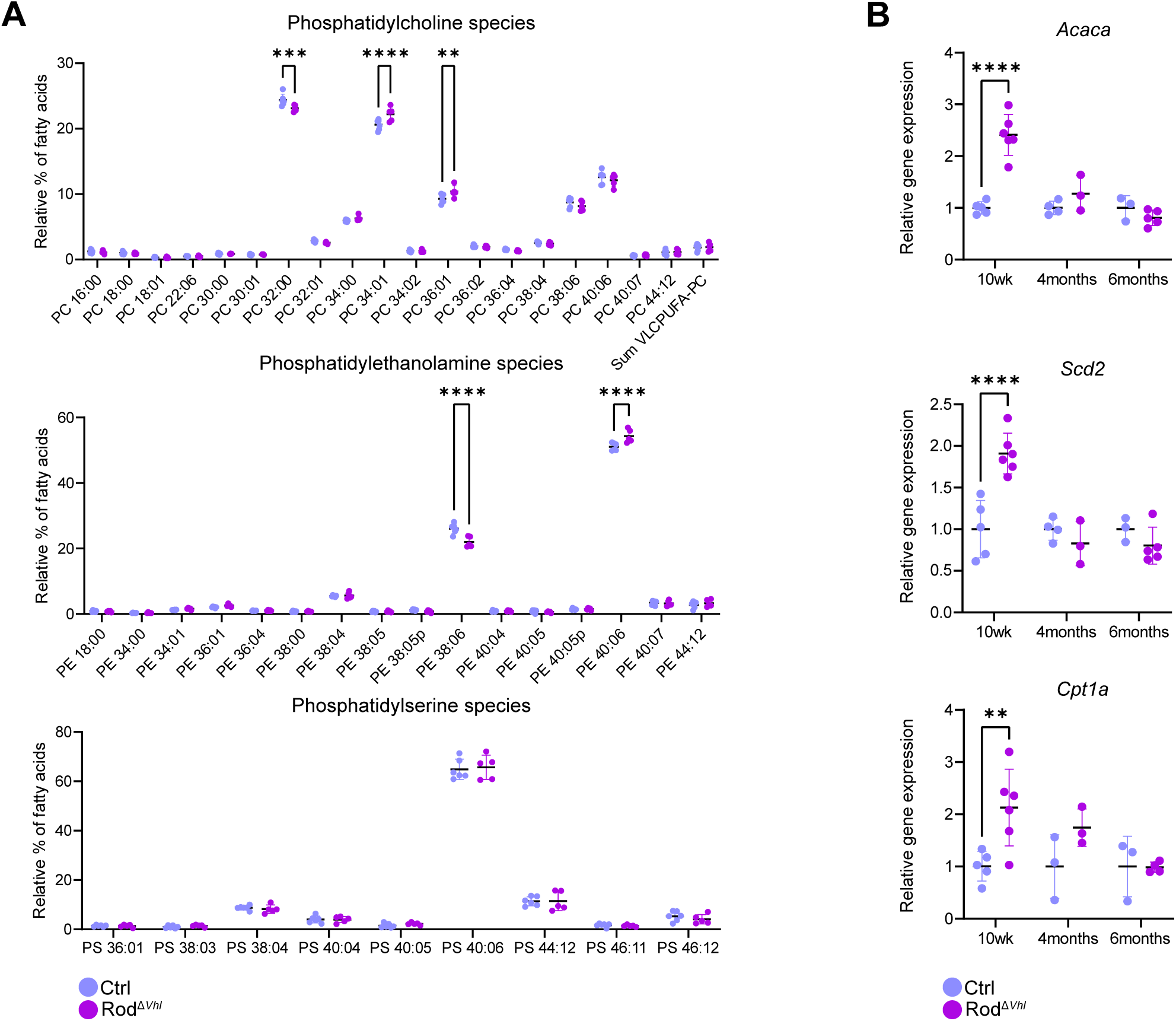
Chronic HIF activity is associated with selective alterations in phospholipid composition in rod outer segments. A, Quantification of PC, PE, and PS species in rod outer segments isolated from retinas at 10 weeks of age (N = 6 samples per genotype; both eyes from 2 mice were pooled per sample). B, qPCR analysis of indicated transcripts in entire retinas of Rod^Δ*Vhl*^ and control mice at 10 weeks, 4 months and 6 months of age. Shown are individual data points (N = 3-6), mean ± SD. *: p ≤ 0.05. **: p ≤ 0.01. ***: p ≤ 0.001. ****: p ≤ 0.0001.

## Discussion

Age-related macular degeneration is a progressive retinal disorder that is caused by multiple factors such as genetic susceptibility, aging and environmental risks [30]. A hallmark of the pathophysiology is the accumulation of extracellular lipid and lipoprotein deposits in the retina, RPE and Bruch’s membrane [31–33] as well as a reduced choroidal blood flow [6, 7, 34] leading to reduced tissue oxygenation [35] and activation of a hypoxic response in cells of the aging retina [11]. Since the outer retina heavily relies on the choroidal circulation for glucose and oxygen, which are essential for maintaining photoreceptor function, we investigated the early metabolic alterations in the retina of a mouse model with chronic HIF activation in rods that mimic the hypoxic response in the aging retina. By characterizing the metabolic signature using a layer-specific multi-omics approach as well as metabolic flux experiments, we aimed to identify potential targetable mechanisms relevant to hypoxia-mediated diseases.

Photoreceptors are extremely vulnerable to oxidative stress due to their high metabolic activity, their continuous exposure to oxygen, as well as the production of ROS during light exposure [36]. In addition, hypoxic conditions, as they occur in the aging eye, can disrupt mitochondrial electron transport, leading to increased electron leakage and the generation of ROS, which can significantly damage the structure of the photoreceptors by oxidizing lipids, proteins and nucleic acids [37]. This puts a high metabolic demand on photoreceptor cells to sustain anabolic production to prevent degeneration [36]. To counteract this, photoreceptors established multiple antioxidizing systems for protection, such as NADPH-dependent pathways, antioxidants including glutathione, and a range of detoxifying enzymes [38]. Although our Rod*^ΔVhl^* mice were raised in normoxic conditions, the chronic activity of HIF was sufficient to lead to an increased oxidative load that was evident in our proteomic and metabolomic datasets, which showed an increase of antioxidant pathways (Figure 1B,C) as well as biochemical indicators of oxidative damage (Figure 1D).

The metabolomics dataset further supported the presence of oxidative stress. Reduced levels of nucleotide-related metabolites suggest impaired nucleotide homeostasis, which is compatible with an oxidative stress state in which nucleotide synthesis, turnover, and recycling may be altered. This interpretation was reinforced by the reduced ratio of hypoxanthine over xanthine. Hypoxanthine is the breakdown product of adenine nucleotides and is a critical intermediate in purine metabolism. Xanthine oxidoreductase converts hypoxanthine to xanthine and subsequently to urate, which is further oxidized to allantoin in rodents [39]. The decreased hypoxanthine/xanthine ratio may reflect reduced nucleotide availability and/or altered purine turnover under oxidative or metabolic stress. The decrease in cAMP and cGMP also supports impaired nucleotide homeostasis, as depletion of the nucleotide pool limits cyclic nucleotide synthesis, while oxidative conditions promote cyclic nucleotide degradation [40]. This metabolic profile of the photoreceptors thus suggests an increased oxidative environment due to chronic HIF activation, which could be detrimental to the structural components of the cell and ultimately result in degeneration that is observed with age in the Rod*^ΔVhl^* mice [13]. However, the cellular response to hypoxia is not limited to changes in the redox status. Activation of HIF can also induce intracellular metabolic pathways that allow cells to adapt to reduced oxygen availability, thereby sustaining their functions and prolonging survival. This became evident when we traced glucose flux in the whole retina of Rod*^ΔVhl^* mice.

Steady-state measurements of retinal glucose uptake, lactate release, and total ^13^C-labeling remained unchanged in the Rod^Δ*Vhl*^ model, indicating that overall glycolytic throughput was preserved in pseudo-hypoxic conditions. This may be explained by the increase of glycogen-related enzymes observed in the Rod^Δ*Vhl*^ photoreceptor layer, which is a well-known metabolic stress response [41]. Photoreceptors might buffer intracellular glucose availability with glycogen metabolism to maintain glycolytic throughput when downstream metabolic pathways are stressed. We cannot completely exclude that stable glucose handling by other retinal cell types masked rod-specific metabolic alterations when measuring whole retinal explants. However, the preserved glycolytic flux combined with increased expression of glycogen metabolism-associated enzymes and the knowledge of the photoreceptors’ intense metabolic activity support the concept that rods activate an early compensatory metabolic response to maintain essential metabolic pathways during hypoxic stress.

While the steady-state glycolytic flux appeared to be preserved, dynamic ^13^C-glucose tracing revealed that chronic HIF activation accelerated the glycolytic flux into downstream branch point pathways (Figure 3E). Glucose enters glycolysis as a six-carbon molecule, glucose-6-phosphate, and is cleaved into two three-carbon intermediates, DHAP and glyceraldehyde 3-phosphate, which are used to generate further metabolites including 3-PG and PEP. The increased ^13^C labelling of these metabolites suggests an acceleration of the early steps of glycolysis, calling for an increased activity or presence of the enzymes involved in preparative glycolysis, which was indeed observed in our earlier work [13]. Since labelling of pyruvate, lactate and TCA cycle intermediates was unchanged, photoreceptors under chronic HIF activation may reroute a part of the three-carbon products derived from glucose while preserving downstream glycolytic and mitochondrial fluxes. A potential explanation is the rate limiting step of pyruvate kinase, which may restrict conversion of PEP to pyruvate, leading to accumulation of upstream glycolytic intermediates that can be diverted into anabolic pathways such as lipid synthesis [42]. Interestingly, the dynamic flux experiments in the RPE showed moderately increased labelling of the different intermediates at early time points in the Rod*^ΔVhl^* tissue, indicating that the RPE cells responded to the altered photoreceptor metabolism by adapting their own internal metabolic pathways.

The increased labelling of DHAP, glycerol-3-P and glycerol suggested a tendency towards increased carbon flux into pathways associated with glycerolipid metabolism, as these metabolites form the three-carbon backbone for phospholipid synthesis [43]. Although steady-state levels of glycerol-3-P in the metabolomics dataset were decreased, this likely reflected rapid turnover rather than reduced production. Glycerol-3-P may be rapidly recycled through the glycerol-3-P shuttle or incorporated into lipid biosynthesis, where it serves as a central precursor for phospholipids and triacylglycerols [44]. Under hypoxic conditions, photoreceptors experience increased oxidative stress, which damages the polyunsaturated phospholipids in the outer segments and increases the need for continuous membrane repair and replacement [45–47]. Redirecting glycolytic intermediates into glycerolipid synthesis may therefore be a metabolic adaptation response to increased oxidative stress, as it generates the necessary components for the renewal of phospholipids to support outer segment disc biogenesis. In parallel, the glycerol-3-P shuttle contributes to cytosolic NAD^+^ regeneration through the interconversion of DHAP and glycerol-3-P, linking glycolytic activity to redox homeostasis [48]. Increased activity of this shuttle in Rod^Δ*Vhl*^ photoreceptors could support both NAD⁺ regeneration and the supply of lipid backbones for outer segment membrane renewal in an attempt to delay degeneration.

This adapted metabolic response toward the fatty acid biosynthesis pathway is also reflected in the lipid composition of the outer segments, where distinct changes in phospholipid species were detected in Rod^Δ*Vhl*^ mice already before the onset of degeneration. Lipidomic analysis revealed a modest increase in monounsaturated PC species (PC34:1 and PC36:1) accompanied by a slight reduction of the fully saturated PC32:0 species, indicating a mild shift toward increased desaturation. This is reflected by a shift in the acyl chain composition of PC from saturated fatty acids to monounsaturated fatty acids (MUFAs). Lipid peroxidation is initiated at double bonds and therefore primarily affects unsaturated fatty acids [49]. Because of this, MUFAs are less prone to peroxidation than polyunsaturated fatty acids, thereby supporting greater resilience to oxidative damage [49]. The shift to MUFA-containing PCs may, therefore, be a protective remodelling strategy of the photoreceptor during chronic oxidative stress. This is consistent with the increased expression of *Scd2*, which is the enzyme that introduces a double bond into saturated fatty acids and converts 16:0 into 16:1 and 18:0 into 18:1. These fatty acids are then incorporated into the PC molecules, increasing the abundance of PC34:1 and PC36:1 potentially at the expense of PC32:0.

Chronic HIF activation also altered the composition of PEs in the outer segments. PE species are enriched in long-chain polyunsaturated fatty acids, such as DHA, which are critical for the outer segment of photoreceptors, for the shape of the disc membranes, and for phototransduction [50]. Our analysis showed that there is a shift from PE38:6 to PE40:6, suggesting increased incorporation of long-chain acyl groups. PE38:6 is highly susceptible to lipid peroxidation due to the high DHA content and its position within the discs [51]. The reduction of this species could reflect the increased turnover of oxidized PE molecules. The increase in PE40:6 suggests upregulation of elongation pathways to maintain the polyunsaturated lipid pool essential for outer segment renewal. Upregulation of *Acaca* gene expression supports this remodelling by increasing malonyl-CoA availability, the key substrate required for elongation of polyunsaturated fatty acids. Enhanced activity of this enzyme would fuel the synthesis of very-long-chain PUFAs incorporated into PE 40:6 and help replenish oxidized PE species during outer segment renewal. While elongation of DHA-rich PEs does not reduce the intrinsic susceptibility of highly unsaturated PE species to peroxidation, it may influence membrane organization and the spatial distribution of reactive sites. On top of this, increased *Cpt1a* expression suggested elevated fatty acid beta-oxidation, which is an important pathway for ATP production during metabolic stress [52]. CPT1A-dependent oxidation also maintains the redox balance and supports mitochondrial function, which could give the photoreceptors more energetic and metabolic flexibility during lipid remodelling [53]. Lastly, the lack of changes in PS supports the selective nature of the observed lipid remodelling in our model, since PS is mostly found on the cytoplasmic leaflet of the cell membrane instead of the disc membranes and thus is less exposed to redox alterations [49]. Therefore, our lipid profiling shows that photoreceptors engage in active remodelling of their phospholipid species content to become more resistant against oxidative stress under chronic HIF activation in an attempt to preserve outer segment integrity and delay degeneration. We hypothesize that these protective pathways, however, ultimately reach exhaustion, as they are energetically very costly and cannot be sustained indefinitely. Lipid remodelling requires a substantial amount of acetyl-CoA, ATP and NADH, which will become increasingly limited under conditions of chronic HIF activation due to metabolic reprogramming [54]. Over time, increased oxidative stress can overwhelm the available antioxidant systems, accelerating lipid peroxidation beyond the compensatory capacities of the cell. This is reflected in our qPCR data, where, in contrast to the early time points, the enzymes involved in fatty acid biosynthesis no longer showed any difference between Rod^Δ*Vhl*^ and control mice at 4 and 6 months, time points of ongoing retinal degeneration. As biosynthesis can no longer keep up with the substantial amount of oxidative damage, membranes become progressively more damaged, and degeneration accelerates. Therefore, prolonged exposure to HIF activation could ultimately contribute to the transition from resilience to cell death.

This compensatory-to-exhaustion hypothesis has important implications for the treatment of retinal degenerative diseases. Many disorders of the outer retina, such as age-related macular degeneration and retinitis pigmentosa, are characterized by chronic metabolic stress, increased oxidative damage, and impaired outer segment renewal [37]. Our findings at the pre-degeneration time point suggest that HIF-induced metabolic adaptation initially supports photoreceptor resilience towards oxidative damage, but the degeneration at later time points indicates that this protective phase will eventually collapse once the energetic and biosynthetic burden becomes unsustainable [13]. This is consistent with previous findings that demonstrated that ablating *Vhl* in rod photoreceptors prolonged photoreceptor survival in a mouse model for retinitis pigmentosa by enhancing glycolysis and improving metabolic support from the RPE [55]. This may be due to the lipid remodelling that is initiated by photoreceptors during chronic HIF activation. Despite these metabolic adaptations, however, HIF stabilization ultimately did not prevent photoreceptor degeneration. Their findings highlight that early HIF-driven metabolic reprogramming can have beneficial effects as long as the compensatory pathways remain intact. Together, these observations emphasize that HIF-driven metabolic remodelling is a double-edged sword, where it is protective in the early phase but may ultimately be insufficient when disease burden exceeds metabolic capacity. These dynamics emphasize the importance of therapeutic timing, as metabolic interventions may only be effective within a limited window before the compensatory capacity is exhausted.

In summary, our work shows that photoreceptors exposed to chronic HIF activation undergo a shift toward an oxidative redox environment, increasing the oxidation of essential cellular components such as nucleotides and lipids. This oxidative pressure triggers the activation of antioxidant pathways, while the transcription factor HIF drives metabolic rerouting towards lipid remodelling, a response that likely helps to sustain outer segment membrane composition and maintain structural integrity under stress conditions. Together, these early adaptive responses highlight how HIF-induced metabolic plasticity initially supports photoreceptor resilience before ultimately reaching its limits as degeneration progresses. Since our data not only reveal metabolic consequences of chronic HIF activation in a model relevant to the aging human retina but also identify cellular alterations before onset of degeneration, they provide potential targets for interfering with disease development. Supporting mechanisms that balance the redox status and normalize metabolism in photoreceptors may have a high therapeutic potential.

## Acknowledgements

We thank the Laboratory Animal Services Center (LASC) of the University of Zurich for animal care and the metabolomics facility of the Functional Genomics Center Zurich for sample processing and data acquisition. This work was supported by the Swiss National Science Foundation (310030_200798) and Hermann Kurz Stiftung.

## Table legends

Supplementary Table S1. **Overview of non-glycolytic and non-oxidative phosphorylation proteins identified in individual retinal layers.** Previously published proteomics data from individual retinal layers, including photoreceptors segment (PS) and outer nuclear layer (ONL), were reanalyzed after exclusion of proteins associated with OXPHOS and glycolysis. Only significantly enriched annotations (*p* ≤ 0.05) are shown with the corresponding abundance ratio.

Supplementary Table S2**. Raw metabolomics data from combined photoreceptor segments and outer nuclear layer samples**. Metabolomics profiling data for the combined photoreceptor segments and outer nuclear layer are presented. For each detected metabolite, retention time (RT, min), mass-to-charge ratio (m/z), fold change (FC), log₂-transformed fold change [log₂(FC)], raw p value, and −log₁₀(p) are reported. In addition, normalized metabolite intensities for all analyzed samples are provided.

